# Serological Profile Of Specific Antibodies Against Dominant Antigens Of SARS-CoV-2 In Chilean COVID-19 Patients.

**DOI:** 10.1101/2021.02.05.429566

**Authors:** K. Cereceda, R. González-Stegmaier, JL. Briones, C. Selman, A. Aguirre, G. Valenzuela-Nieto, C. Caglevic, R. Gazitua, A. Rojas-Fernandez, F. Villarroel-Espíndola

## Abstract

Coronavirus disease 2019 (COVID-19) is caused by SARS-CoV-2 and has been a pandemic since March 2020. Currently, the virus has infected more than 50 million people worldwide and more than half a million in Chile. For many coronaviruses, Spike (S) and Nucleocapsid (N) proteins are described as major antigenic molecules, inducing seroconversion and production of neutralizing antibodies. In this work, we evaluated the presence in serum of IgM, IgA and IgG antibodies against N and S proteins of SARS-CoV-2 using western blot, and developed an ELISA test for the qualitative characterization of COVID-19 patients. Patients with an active infection or who have recovered from COVID-19 showed specific immunoblotting patterns for the recombinants S protein and its domains S1 and S2, as well as for the N protein of SARS-CoV-2. Anti-N antibodies were more frequently detected than anti-S or anti-S1-RBD antibodies. People who were never exposed to SARS-CoV-2 did not show reactivity. Finally, indirect ELISA assays using N and S1-RBD proteins, alone or in combination, were established with variable sensitivity and specificity depending on the antigen bound to the solid phase. Overall, Spike showed higher specificity than the nucleocapsid, and comparable sensitivity for both antigens. Both approaches confirmed the seroconversion after infection and allowed us to implement the analysis of antibodies in blood for research purposes in a local facility.

## INTRODUCTION

Coronavirus disease 2019 (COVID-19) is caused by the severe acute respiratory syndrome coronavirus 2 (SARS-CoV-2). The virus emerged in December 2019, spreading quickly, and was subsequently declared a worldwide pandemic by the World Health Organization in March 2020 (1–3). SARS-CoV-2 virus is a beta coronavirus and its structure includes four major structural proteins and several accessories proteins (Orf, open reading frame) Orf1ab, Orf3a, Orf6, Orf7a, Orf8 and Orf10. Regarding the structural components, M protein is a membrane (M) protein which participates in virus assembly and stabilizes the Nucleocapside-RNA interaction (4); the E protein is part of the envelope (E) and is involved in virus particle replication including assembly and budding; the Nucleocapsid (N) protein is responsible for binding to the single strain RNA (ssRNA) and plays a role in virus replication; and finally, the Spike (S) protein has a specific receptor binding domain (RBD) that interacts with the receptor in the host cells allowing the virus entry through its interaction with ACE2 (5). For many coronaviruses, S and N proteins have been described as major antigenic molecules, which induce, in most of people, seroconversion and production of a neutralizing antibody. SARS-CoV and SARS-CoV-2 share a 76% amino acid identity across the genome (6,7). Several nonsynonymous mutations were described in the S protein during the SARS-CoV epidemic progression (8,9), and it is already happening for the SARS-CoV-2 pandemic (7,10), suggesting that Spike may be a unique antigen allowing us to distinguish between coronavirus strains. In contrast, the N gene is more conserved and stable, with 90% amino acid homology and fewer mutations over time (11–14), and it is considered as one of the most immunogenic antigens in SARS-CoV and SARS–CoV-2 (15–17). In general, antibody responses generated against the S and the N proteins have shown to protect mice from SARS-CoV infection and have been detected in SARS-CoV and SARS-CoV-2-infected patients (18–23). Regarding coronavirus-induced antibodies, the S protein is highly immunogenic to induce antibody responses in SARS patients and is a potent inducer of neutralizing antibodies against SARS-CoV in immunized animals (24–26). This protein has been extensively used as a major antigen for developing diagnostic tools and as a major immunogen for developing SARS vaccines. Interestingly, murine polyclonal antibodies developed against S protein from SARS-CoV showed an inhibitory effect on SARS-CoV-2 S-mediated entry into cells, indicating the presence of dominant and conserved epitopes which can allow cross-reactivity and cross-neutralizing antibodies (5,27). On the other hand, N proteins of many coronaviruses are highly immunogenic and these are expressed abundantly during infection (17,28). The middle or C-terminal region of the SARS-CoV N protein is known for eliciting antibodies against SARS-CoV during the immune response (15,29,30). High levels of IgG antibodies against N have been detected in sera from SARS patients (31), and the N protein is a representative antigen for the T-cell response in a vaccine setting, inducing SARS-specific T-cell proliferation and cytotoxic activity (17,32). IgM and IgG antibodies against the N protein of SARS-CoV-2 have been frequently detected in COVID-19 patients and those have shown to be more sensitive than spike protein antibodies for detecting early infection (33,34). So far, the measurement of serum antibodies has been reported as crucial data for understanding aspects of diagnosis, prognosis and public health decisions regarding SARS-CoV-2 infection (35). In this work we evaluated the presence in serum of IgM, IgA and IgG antibodies against selected antigens from SARS-CoV-2 N and S proteins using western blot for selecting antigens based on its nonconform ational immunogenicity based on its primary structure. Collected serum from patients with an active infection or recovered from COVID-19 showed a specific immunoblotting pattern for the recombinant SARS-CoV-2 S protein and its domains S1 and S2, as well as for the N protein; however, the antibodies against the N protein were more frequently detected than antibodies anti-S or anti-S1 RBD. No reactivity by western blot was observed using serum from people never exposed to SARS-CoV-2. The three isotypes IgM, IgA and IgG antibodies against S or N proteins were individually detected and they showed dramatic differences at the level of abundance, temporality and specificity. In addition, we developed and validated an indirect ELISA assay using N and S1-RBD proteins, alone or in combination, for the qualitative detection of antibodies in plasma or serum from COVID19 patients. The sensitivity (Se) and specificity (Sp) of the ELISA assay was 100% and 71% for the N protein alone, 99% and 85% for the S1-RBD alone, and 100% and 55% respectively for both combined. Both approaches confirmed the seroconversion after infection and allowed us to implement the analysis of antibodies in blood for research purpose in a local facility.

## MATERIALS AND METHODS

### Patients and samples

All samples related to COVID-19 were collected between April 4^th^ and May 15^th^ of 2020. Recovered patients and those with an active infection with SARS-CoV-2 were invited to participate and signed a consent letter at the moment of enrollment to the clinical trial NCT04384588. Briefly, patients with severity criteria had any of the following conditions: dyspnea and or respiratory rate >=30 per min and or saturation <= 93% with fraction of inspired oxygen 21% and/or ratio of partial pressure arterial oxygen and fraction of inspired oxygen (PaFi) <300 and/or lung images showing worsening in 24-48 hours. Patients were recruited and consented at the moment of hospitalization and classified as critically ill participants for this work. As exclusion criteria any known allergy to plasma, a severe multiple organic failure, an active intra brain hemorrhage, disseminated intravascular coagulation with blood products requirement, patient with an adult respiratory distress longer than 10 days, patients with active cancer and life expectancy shorter than 12 months according with medical criteria was considered. Recovered people from COVID-19 were invited to be volunteers for convalescent plasma donation through the clinical trial indicated above; they consented and signed a consent letter at the apheresis day. Inclusion criteria for convalescent patient considered men and women previously confirmed with COVID-19 by PCR test and 21 or more days after symptoms had finished. Any convalescent participant who had showed a second positive PCR test within the third week of recovering was excluded for this research. The never exposed to SARS-CoV-2 control sample was obtained from a single blood drawn in April 2020 from asymptomatic healthy volunteers with a negative SARS-CoV-2 PCR result.

All samples were recovered from the institutional Biobank at Fundación Arturo López Pérez Cancer Center. Briefly, the blood sample was collected in a tube containing potassium EDTA, then centrifuged at 1500 xg for 15 minutes within 60 minutes of blood draw, aliquoted and kept frozen at −80°C until processing for investigation. A summary of total cases, clinical and demographic information is included in supplementary data (Tables S1 and S2).

### Recombinant proteins and secondary antibodies

Recombinant SARS-CoV-2 Nucleocapsid Protein with C-terminal His-tag (#230-30164-100); Recombinant SARS-CoV-2, S1 Subunit Protein (RBD) with C-terminal His-tag (#230-30162-100); Recombinant SARS-CoV-2 Spike Protein, S1 Subunit; Recombinant SARS-CoV-2 Spike Protein (#230-01101-100), S2 Subunit (#230-01103-100) were purchased from Raybiotech (Peachtree Corners, GA, USA). Peroxidase-conjugate Donkey anti Human-IgG (#709-035-098), Peroxidase-conjugate Donkey anti Human-IgM (#709-035-073) and Peroxidase-conjugate Donkey anti Human-IgA (#109-035-011) and Cy3-conjugated Donkey anti Human IgG (#709165098) were purchased from Jackson ImmunoResearch (West Grove, PA, USA). 4’,6-Diamidino-2-phenylindole dihydrochloride (DAPI) (NBP2-31156-1mg) probe was purchased from Novus Biologicals (Centennial, CO, USA)

### SDS-PAGE

500 ng for each recombinant SARS-CoV-2 protein was prepared for electrophoresis using 4x Laemmli Sample Buffer (#1610747, Biorad Laboratories, Hercules, CA, USA) and incubated for 5 minutes at 95°C and then separated by 12% SDS-PAGE. A broad-range protein marker (10-250 KDa, Kaleidoscope # 1610375, Biorad) was included. After the electrophoretic separation, the gels were either stained with One-Protein Blue gel stain (#21003-1L, Bioticum) or transferred onto PVDF membranes for immunodetection using TurboBlot system (Biorad).

### Western Blot

The membranes were blocked for 30 minutes at room temperature with 5% bovine serum albumin (BSA) in TBS-T (100mM Tris-buffered and 0.1% Tween-20). Later, the blocking solution was removed and individual membranes were separately incubated overnight at 4°C in a rocking platform with each serum sample diluted at 1:500 in blocking solution. The next day, all solutions were removed and the membranes washed three times with a TBS-T solution for 5 minutes each time. For the immunodetection, the membranes were separately incubated with either HRP-conjugated secondary antibody anti-human IgG, anti-human IgA or anti-human IgM diluted at 1:20,000 (0.04 μg/ml) in blocking solution. Secondary incubation was performed at room temperature for 1 hour and agitation (200 rpm). The antigen-antibody interaction was developed with Clarity Western ECL Substrate (#1705060, Biorad) and detected in an Omega Lum C detector. Pixels were quantified using Fiji Software (v.2.0.0, NIH) and exposition time was used as reference.

### Immunofluorescence

HeLa cells (ATCC^®^ CCL-2) were transient transfected with a mammalian expression vector expressing a Spike-GFPSpark tag codon optimized fusion SinoBiological VG40589-ACGLN in 10cm plates, 24 h after transfection ~ 8000 cells per well were deposited into 96 well optical plate (Themofisher), after 24 h incubation cells were washed with PBS (phosphate buffered saline solution) 1X 3 times and fixed with 4% paraformaldehyde at room temperature for 30 min. After fixation, cells were washed with PBS 1X and permeabilized in PBS 1X 0.2% Triton X-100 and further washed 3 times in PBS 1X and store at 4°C (36). Random serum samples from COVID-19 recovered patients were assessed for staining Spike protein by immunofluorescence. Cells were incubated for 30 minutes at room temperature with blocking solution containing 0.5% BSA, 0.1% non-immune Donkey serum (#017-000-121, Jackson ImmunoResearch), and 0.1% Tween-20. Cells were then incubated for 30 minutes at 37°C with either control serum from healthy patients or with serum from COVID-19 patients diluted at 1:100 in blocking buffer. Cells were washed with PBS 1X and then incubated with Cy3-conjugated anti-Human IgG diluted at 1:400 (3.75 μg/ml) in blocking solution for 30 minutes at 37°C. For nuclei staining, cells were washed with PBS 1X and incubated with 0.2 μg/ml DAPI for 15 minutes at room temperature. Then, images were acquired using a Cytation 5 Cell Imaging Multi-Mode Reader (BioTek, Winooski, VT, USA)

### Indirect ELISA for IgG against S1-RBD SARS-CoV-2

The serum SARS-CoV-2 antibodies were detected using an in-house indirect ELISA test which was developed and validated in the Translational Medicine Laboratory of Foundation Arturo López Pérez Cancer Center in Chile. Briefly, microplates 96 well MaxiSorp^™^ (#439454, NUNC Thermo Sci) were coated overnight at 4°C with 50 ng of each antigen alone (SARS-CoV-2 S1-RBD and N protein) in 0.1M carbonate buffer pH9, then the coating solution was removed and each well was washed once with cold phosphate PBS 1X. Uncoated surfaces were blocked for 4 hours at room temperature using 400μl of a blocking solution containing 5% skim milk in PBS 1X pH7 and supplemented with 0.1% Bovine serum albumin (BSA), 0.1% non-immune donkey serum (#017-000-121, Jackson ImmunoResearch), and 0.1% Tween-20. After blocking, the solution was eliminated and plates were air dried to be stored frozen at −20°C. Frozen plates were stable for up to 45 days based on the measurement of inter-well variation coefficient of the absorbance of a reference material and blank without sample. Before testing unknown samples, plates were thawed at room temperature and the excess blocking solution was removed using 200μl of washing solution containing 0.1% Tween-20 in PBS 1X with an automatic microplate washer and kept at room temperature for seeding the samples. Serum samples, controls and calibrator were diluted fresh at 1:320 using as dilutor a solution containing 0.1% BSA in PBS 1X and supplemented with 0.1% non-immune donkey serum and 0.1% Tween-20. 100μl of each dilution was seeded in duplicate and incubated for 1 hour at room temperature (20 ± 2°C). Using an automatic microplate washer, all liquids were removed and the plate rinsed 5 times (2 sec shaking and 10 sec soaking) with 250ul of washing solution each time. Later, each well was incubated at 20°C (±1°C) for 1h with 100 μL of a 20 ng/ml solution of the specific peroxidase-conjugated affiniPure anti-Human isotype antibody, anti-IgG (Fcβ fragment specific) (#709-035-098, Jackson Immuno-Research), anti-IgA (Frα fragment specific) (#109-035-011, Jackson Immuno-Research) and anti-IgM (μ chain) (#709-035-073, Jackson Immuno-Research). Finally, the plates were rinsed as mentioned above and the reaction developed using a 3,3’,5,5’-tetramethyl-benzidine substrate (TMB) (#T0440, Sigma-Aldrich) for 15 minutes at room temperature and light-protected. The reaction was stopped with 2M of sulfuric acid and absorbance measured at 450 nm in a Cytation5^®^ plate reader (BioTek, Winooski, VT, USA). The performance of blank, controls, and calibrator were supervised based on absorbance <10% CV. The interpretation of the results of the ELISA test allowed two approximations: a qualitative analysis based on the absorbance ratio at 405nm between the unknown sample and a calibrator (known standard positive). Any samples with an absorbance ratio equal or higher than 1.1 were considered positive; negative samples consisted of any with a ratio below 0.9; and samples were considered unknown or non-classifiable when the ratio was between 0.9 and 1.1. A semi-quantitative analysis is allowed based on the continuous value of the calculated ratio for each unknown sample.

Previously tittered sera using an IVD ELISA test (Euroimmun) were used as an internal positive control (ratio above 2.5, 7.5%CV), as a calibrator (ratio 1.0±0.05, <5%CV) and as an internal negative control (ratio below 0.8, 5%CV) according to the manufacture’s indications. <

### Statistical analyses

Statistical analysis was performed using GraphPad Prism software, version 8.0. A preliminary Pearson’s correlation coefficient was used to assess the relationship between absorbance at the dilution 1:320 and the measured ratio by ELISA. Parameters of sensitivity (Se), specificity (Sp), positive predictive value (PPV) and negative predictive value (NPV) were estimated based on the accuracy of the assignment of positive and negative known samples. The concordance coefficient and the percentage of agreement were calculated against an IVD ELISA test and also, a rapid lateral flow assay. The Mann-Whitney test was used to analyze differences in mean values between groups. All values are depicted as mean and standard error. One-way ANOVA and Tukey’s multiple comparisons test were performed for statistical analysis between various variables. The critical value for statistical significance was established as p<0.05.

## RESULTS AND DISCUSSION

### Detection of IgG antibodies against SARS-CoV-2

To determine the presence of specific antibodies against SARS-CoV-2 in serum, 32 samples collected between April and May 2020 in Santiago, Chile were assessed by western blot against recombinant S1 and S2 domains of Spike (S) protein and the whole Nucleocapsid protein (N) of SARS-CoV-2. As shown in figure 1, most of the serum samples from Chilean COVID-19 patients contained specific antibodies and showed strong reactivity. All detected bands for each recombinant were at the expected molecular weight (S1=75kDa; S2=58kDa, and N=57kDa) confirming a polyclonal production of IgG against SARS-CoV-2 (Fig1A). Individuals within 10 days after PCR positive diagnosis of COVID-19 were characterized by a minimal reactivity of IgG against Spike (S1 or S2 domains) where only 2 out of 10 showed positive bands; however, antibodies against the N protein were clearly present in 4 of 10 (Fig 1A). On the contrary, 100% of convalescent samples (12 of 12) from COVID-19 recovered people with 28 or more days after symptoms offset showed IgG antibodies against Spike with a dominant signal for S2 domain, and the presence of anti-N IgG with comparable intensity to those detected during active disease (Fig 1A). In general, the antibodies against the N protein were shown to be more reactive, and a semi-quantitative analysis after exposure to time normalization showed that anti-N IgG may dominate in abundance independently of the stage of disease (Fig1B). In fact, N-protein has been described as the more abundant viral protein by several groups (37–39) and extremely immunogenic (16,40). Regarding the presence of antibodies within the first days after infection, a quick seroconversion has been reported in some cases (33,34,41,42) and suggested an increased seroconversion rate from week 1 (25.9%) to ≥4 (96.5%) post symptoms onset (42). Recently, a clinical study which was attempted using convalescent plasma in severe cases of COVID-19 found that 53 of 66 of participants (less than 10 days of symptoms onset) already had antibodies against SARS-CoV-2 at baseline (41). So far, those reports confirm our findings by western blot.

**Fig 1.**
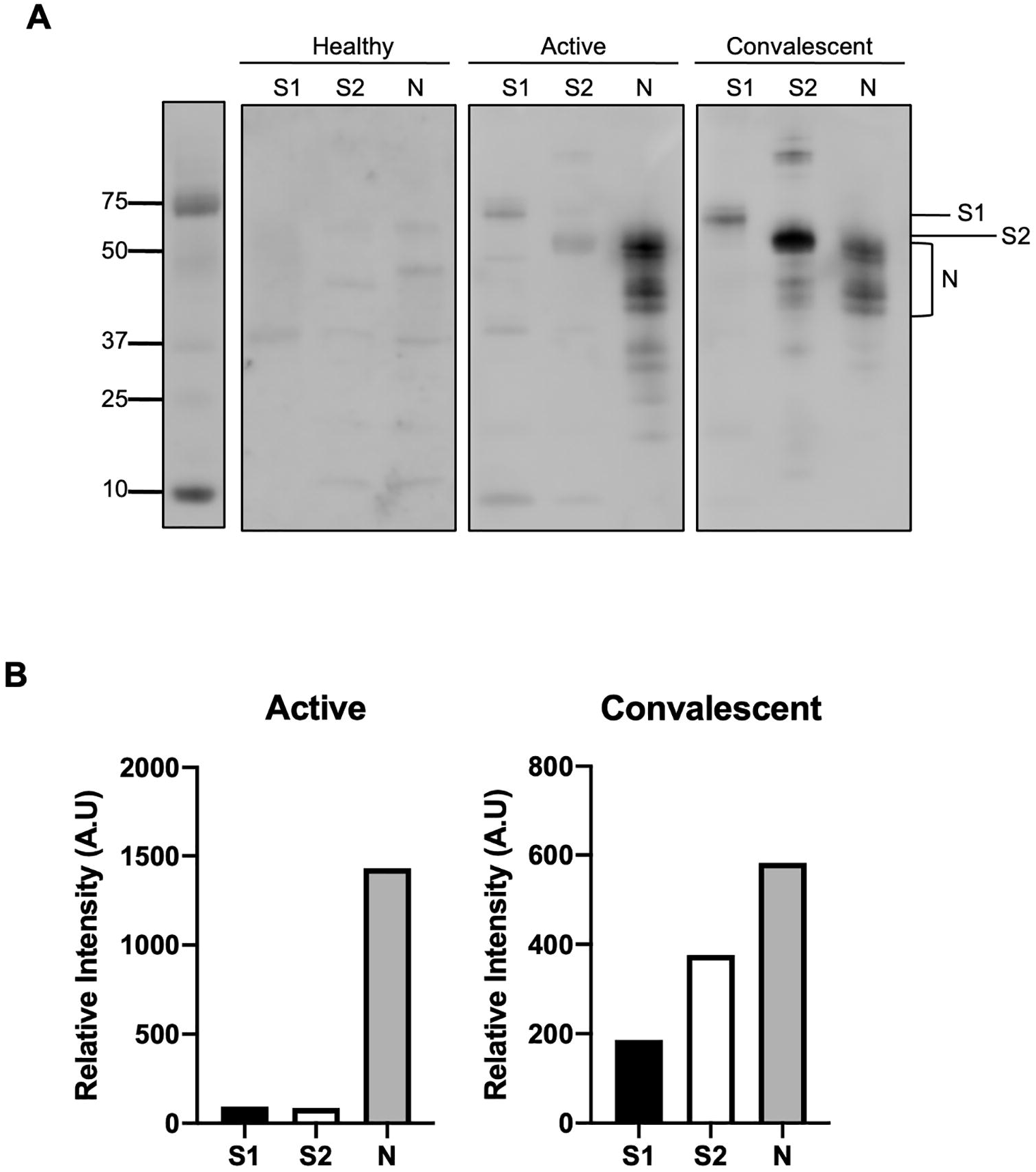
Immunodetection of serum IgG against Spike-subunits (S1 & S2) and N-protein in patients with COVID-19 active and convalescent. (A) Western blot analysis of IgG anti S1 (75 KDa), S2 (58 KDa) and N (47 KDa) protein in serum from an active and convalescent COVID-19 patient. (B) Densitometric analysis of Western Blot in A. Exposure time was used to normalize the signal detection for intensity semi-quantification. (C) Mean ± SEM are indicated for each group. Blot images and relative abundance graphs are representative of ten independent cases. *Und* undetectable.

### Functional isotypes anti-Spike of SARS-CoV-2

Previously, the FDA approved the use of convalescent plasma as an emergency strategy in COVID-19 patients (FDA & Kadlec, 2020) and recommended measuring IgG as standard. To generate preliminary data of seroconversion and the feasibility of using Chilean convalescent plasma, in addition to the IgG antibodies, the presence of IgM and IgA against Spike was analyzed in the convalescent group (Fig2). All cases (12 of 12) showed IgA and IgM reactivity after 35±7 days after symptoms offset (Fig 2A). After normalizing the exposure time, reactive bands were quantified and showed a prevalence of IgG over IgM and IgA, with specificity Spike, in particular against the S2-domain (Fig 2A and 2B), and very low detection of S1-RBD, which is the target to neutralize SARS-CoV-2 infection (Fig 1A and 2A). To understand the neutralizing potential of antibodies in plasma, we assessed the presence of antibodies anti-RBD in active COVID-19 samples using single blotted strips. After individual western blot analysis, only one case showed a clear IgG immune-reactive signal anti-RBD (less than 10 sec), and 9 out of 10 required at least 120 sec to generate a successful image (Fig 2C).

**Fig 2.**
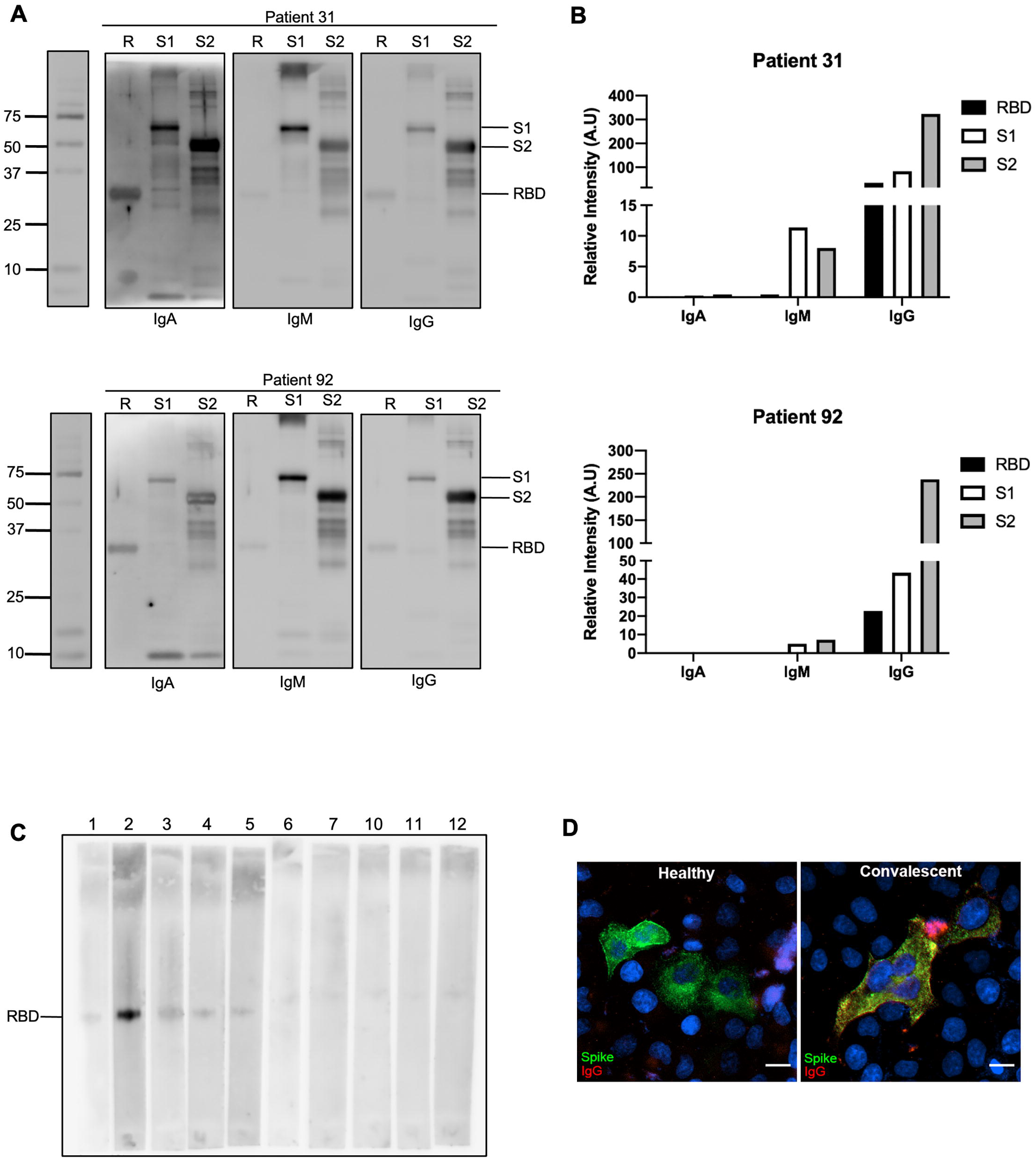
Immunodetection of serum antibodies IgA, IgM and IgG against −RBD, −S1 and −S2 subunits (Spike) in patients recovered from COVID-19. (A) Western blot analysis of IgA, IgM and IgG against S1 (75 KDa), S2 (58 KDa) Spike subunits and RBD (30 KDa) in serum from 2 different patients recovered from COVID-19. (B) Densitometric analysis of Western Blots in A. Exposure time was used as reference for intensity determination. (C) Western blot analysis of anti-RBD IgG (30 KDa) in serum from 10 different active COVID-19 patients. (D) HeLa cells expressing GFP-Spike (Green) were used to determinate anti-Spike IgG (Red) in serum from a patient recovered from COVID-19 compare to a healthy control serum. Merge (Yellow) images are shown. Nuclei were stained with DAPI (blue). Scale bar, 10 μm. Blot images are representative of ten independent cases.

Western blot assays are technically limited to measure antibodies with affinity and binding ability mostly to lineal and sequential epitopes, for that reason we probed the antibodies binding against the full Spike protein in its native 3D conformation. HeLa cells overexpressing the whole recombinant EGFP-Spike of SARS-CoV-2 were incubated in the presence of conditioned medium with each previous tested serum sample. Female healthy control samples were used to define background in case of any anti-HLA cross-reaction, and male samples as negative control as well. As shown in figure 2D, healthy donors did not have antibodies able to bind specifically EGFP-Spike; however, all samples from convalescent people (12 out 12) showed a total co-localization indicated by the yellow pseudo-colored overlap between the green-fluorescent protein of Spike and the alexa-594 emission from the secondary detection of the bound IgG. This result suggests the presence of specific IgG antibodies against structural epitopes present in S1-RBD of SARS-CoV-2 which may have a neutralizing capacity.

### In-house ELISA anti-relevant COVID-19 antigens

Since we were able to detect antibodies against Spike and N protein in its denaturized and native 3D conformation, we decided to develop an indirect ELISA test for testing samples with research purposes. Three in-house ELISA tests to measure antibodies anti-N, anti-S1-RBD or combining both antigens were established (Fig 3A). When the antigens were exposed in their native conformation, most of the previously analyzed samples showed positive results within a range of dilutions between 1:100 and 1:400 (data not shown). The in-house test was defined to be performed at the dilution 1:320 as the FDA recommended for screening and selecting convalescent plasma donors. A small group of 9 independent cases not analyzed previously by western were included. All convalescent cases had antibodies against any of the two major structural proteins of SARS-CoV-2 and the relative amount for each immunoglobulin was statistically higher than in healthy non-COVID-19 people (n=15, p=0.01-0.001). When samples were segregated as positive or negative, they showed 2 of 9, 2 of 9 and 4 of 9 were negative for IgG, IgM and IgA, anti-RBD respectively; 5 of 9, 6 of 9 and 5 of 9 were negative for IgG, IgM and IgA, anti-N respectively; and 2 of 9, 2 of 9, and 0 of 9 were negative for IgG, IgM and IgA, against both antigens respectively.

**Fig 3.**
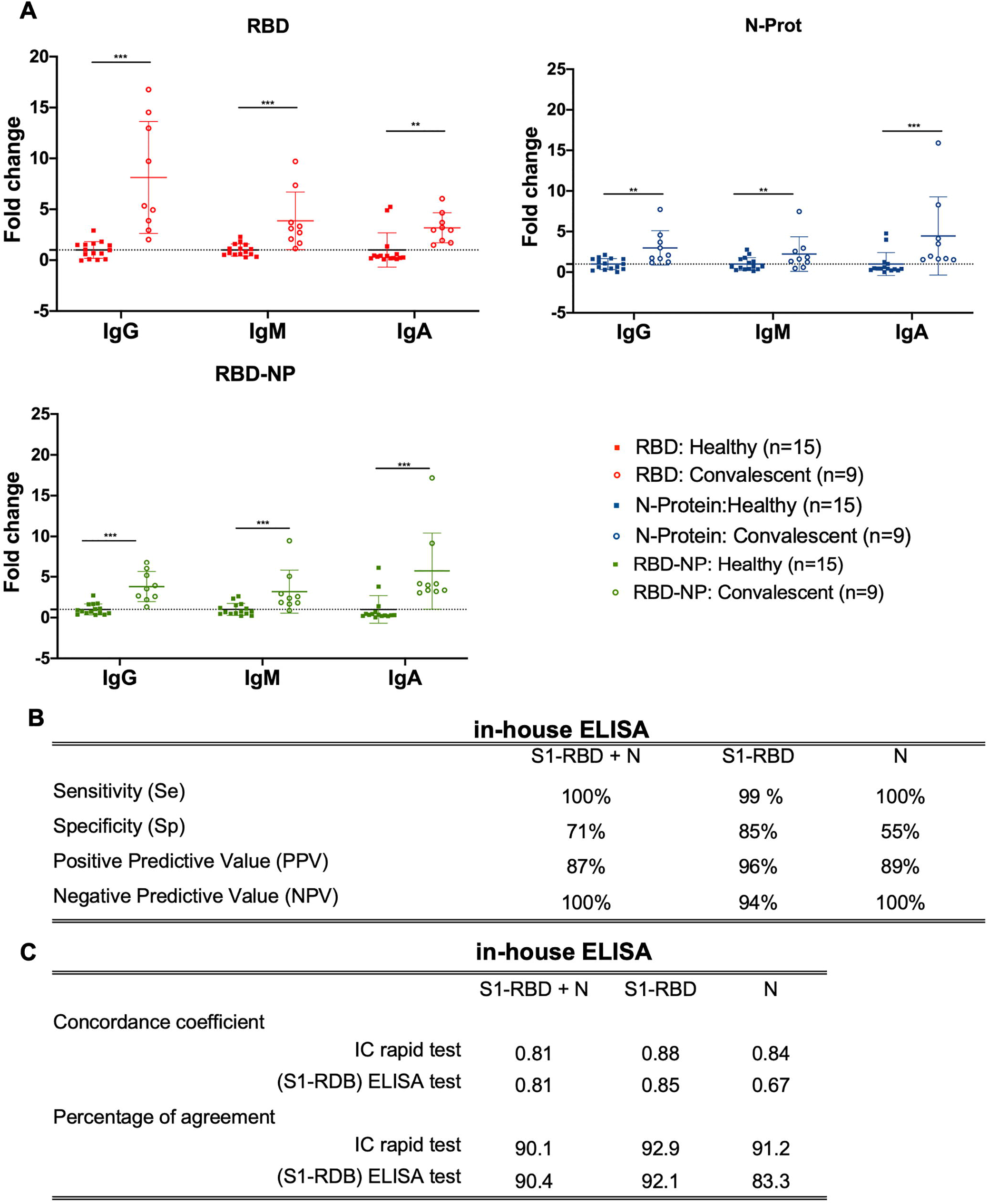
ELISA determination of RBD, N protein and RBD-N protein specific IgG, IgM and IgA antibodies in serum from patients recovered from COVID-19. (A) Data distribution of Convalescent serum results compare to a healthy group. **P<0.01, ***P<0.001. Data are mean with SD. (B), (C) IgG ELISA validation (n=60 cases) previously tested by an IVD ELISA assay using S1-RBD as target (Euroimmun). N= N protein. IC= Immuno-chromatography.

A larger group was used for IgG ELISA validation and it included 68 cases previously tested by an IVD ELISA assay using S1-RBD as target (Euroimmun). When IgG was measured the estimated sensitivity (Se) and specificity (Sp) was 100% and 50% for the N protein alone, 99% and 85% for the S1-RBD alone, and 100% and 71% for both combined (Fig 3B). The positive and negative predictive values reached 89% and 100% for the N-protein, 96% and 94% for the S1-RBD alone, and 87% and 100% for the combined antigens respectively. Sequence analysis of N protein between SARS-CoV-2 and other coronaviruses such as SARS-CoV, MERS-CoV, HKU1 and OC43 revealed a high homology (14,44), suggesting a high degree of cross reactivity induced by previous infection with other coronaviruses. That information may explain the poor performance observed for N protein which showed the highest observed number of false positives within the healthy control group (data not shown). To increase the level of validity, we compared the performance of our ELISA IgG-version against orthogonal IVD tests, and the concordance coefficient and percentage of agreement were calculated. For IgG and only S1-RBD, it showed a coefficient of 0.85 and 0.88, and agreement of 92.1% and 92.9% comparing against an ELISA and a lateral flow assay respectively (Fig 3C). Overall, our in-house ELISA significantly segregated people never exposed to COVID-19 patients, and its versatility allowed the qualitative estimation of three target antibody isotypes (IgA, IgM and IgG) using S1-RBD or N proteins.

For an accurate interpretation of our results, some biochemical subjects needed to be considered. The heterogeneity by immunoblotting may reflect the presence of antibodies against lineal epitopes avoiding the detection of structural epitopes; however, differential reactivity could be observed because the recombinant S1 and S2 peptides were produced in a prokaryotic system and lack of glycosylation, which has been reported as a major immunogenic determinant in coronavirus infection (45). Even though the N protein was produced in HEK293 cells, its observed immunodetection pattern was also variable, reflecting all different posttranslational modifications described for that protein (46,47), as well as the heterogeneity of the IgG clones present in each sample. We understood the limitations of our work in the number of cases analyzed and the experimental strategy used; however, it provides information for some laboratories with limited resources to apply, exceptionally, basic research tools for monitoring COVID-19 cases during a pandemic.

## ETHICS STATEMENT

This study is part of the clinical trial NCT04384588 (clinicaltrials.gov), and met all the required inclusion/exclusion criteria of the mentioned trial. This research and the trial were reviewed and approved by the Institutional Scientific and Ethical Committee of Instituto Oncológico Fundación Arturo López Pérez (Santiago, Chile) and complied all regulatory indications from our Institutional Medical Direction, together with all the legal and ethical requirements of Chilean law.

## AUTHOR CONTRIBUTIONS

Conceptualization: FVE

Methodology: KC, RGS, FVE

Validation: RGS, FVE, CS

Formal Analysis: KC, RGS, FVE

Investigation: KC, RGS, FVE

Resources: RG, JLB, CS, CC, AA, ARF, GVN, JLB, CS

Data Curation: KC, FVE

Writing-Original Draft: KC, RGS, FVE

Writing-Review & Editing: KC, RGS, RG, CC, ARF, FVE

Visualization: KC, RGS, FVE

Supervision: FVE, RGS

Project Administration: FVE

Funding Acquisition: FVE, RG

All authors have read and approved the final manuscript

## LIMITATIONS OF THIS RESEARCH

We recognize the limited number of cases studied in this research, and after analyzing 99 serum samples from people infected with SARS-CoV-2 (recovered and actively sick) we cannot consider our result as representative for a larger population. Our data support the idea of using tools from basic science to determine the presence and absence of anti-SARS-CoV-2 antibodies.

## ACKNOWLEDGMENTS

This study was supported by “Fondo de Adopción tecnológica SIEmpre” sponsored by SOFOFA (Sociedad de Fomento Fabril), CPC Chile (Confederación de la Producción y de Comercio) and Ministerio de Ciencia y Tecnología, Conocimiento e Innovación, Chile. Additionally, ELISA test design was partially supported by CONICYT/FONDECYT Postdoctoral Grant 3170356 (RGS). We also thank Gregory Dunne for the English review assistance. We thank Yanara Delgado and Teresa Alarcon from the Biobank unit.

