## Supplementary figures and images for "Serological Profile Of Specific Antibodies Against Dominant Antigens Of SARS-CoV-2 In Chilean COVID-19 Patients."

### Supplemental Figure 1

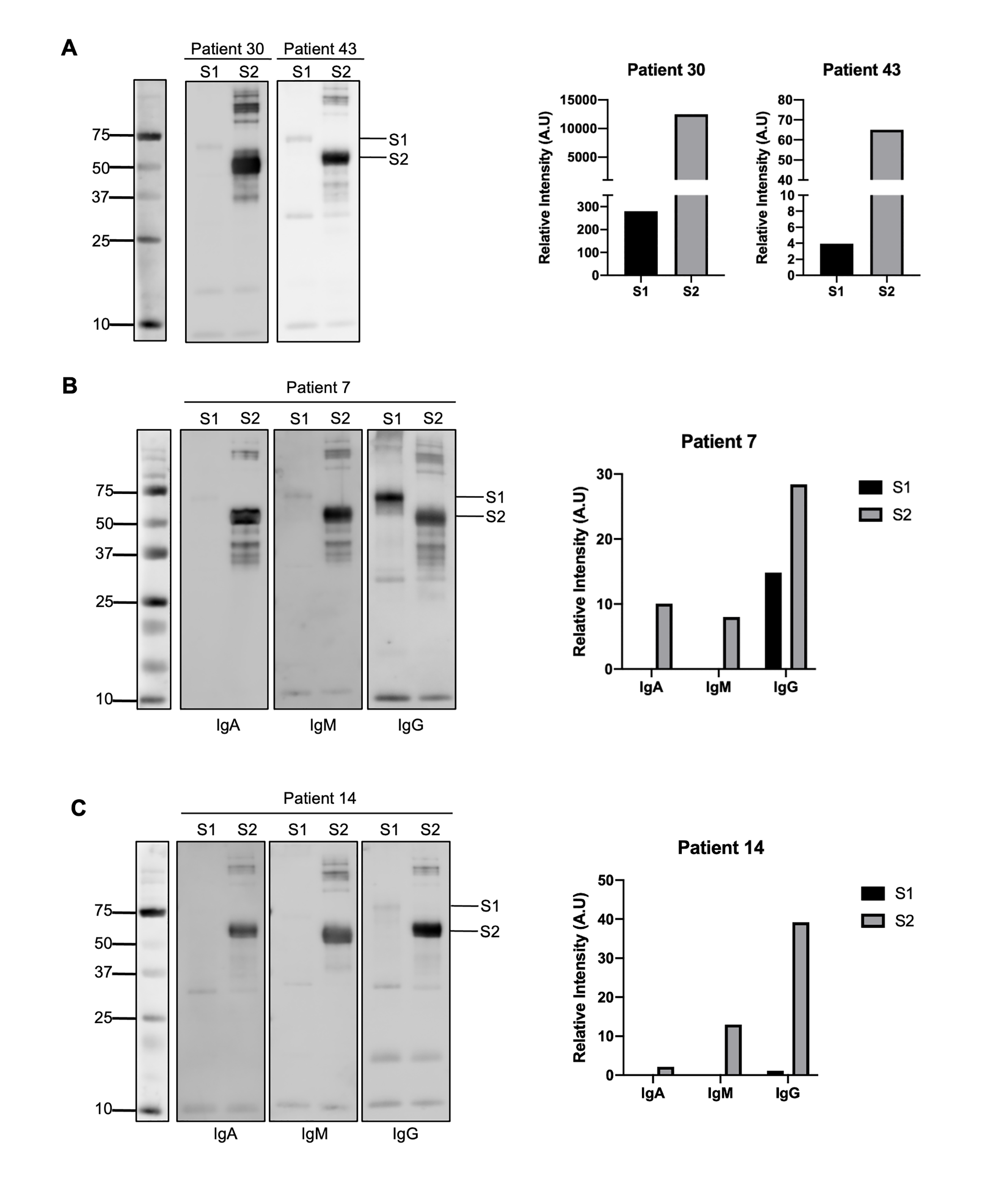
