## Supplemental Table 1 for "Serological Profile Of Specific Antibodies Against Dominant Antigens Of SARS-CoV-2 In Chilean COVID-19 Patients."

**S1 Table.** Demographic and clinical characteristics of the sample cohort used for western bloting

| ***Western blot analysis*** | **Convalescent** | **Active** | **Healthy** |
| --- | --- | --- | --- |
| Total cases | 12 | 10 | 10 |
| Age range (years) | 27-57 | 47-69 | 28-53 |
| Women | 2 | 1 | 6 |
| Men | 10 | 9 | 4 |
| Sample collection |  |  |  |
| Days after symptoms onset | NA | 04-sept | NA |
| Days after symptoms offset | 21-42 | NA | NA |
| Symptomatology during COVID-19 | Mild to Moderate | Severe | NA |
| Symptomatology at collection | Asymptomatic to Mild | Critically ill | Asymtomatic |
| PCR test | Positive | Positive | Negative |
