## Supplemental Table 2 for "Serological Profile Of Specific Antibodies Against Dominant Antigens Of SARS-CoV-2 In Chilean COVID-19 Patients."

**S2 Table.** Demographic and clinical characteristics of the sample cohort used for in-house ELISA development and validation.

| ***In-house ELISA development*** | **Convalescent** | **Healthy** |
| --- | --- | --- |
| Total cases | 9 | 15 |
| Age Range (years) | 24-56 | 28-53 |
| Women | 1 | 4 |
| Men | 8 | 11 |
| Sample collection |  |  |
| Days after symptoms offset | 21-28 | NA |
| Symptomatology during COVID-19 | Mild to Moderate | NA |
| Symptomatology at collection | Asymptomatic to Mild | Asymtomatic |
| PCR test | Positive (100%) | Negative (100%) |
| ***in-house ELISA validation*** | **Convalescent** | **Healthy** |
| Total cases | 68 | 25 |
| Age Range (years) | 23-60 | 28-53 |
| Women | 17 | 12 |
| Men | 51 | 13 |
| Sample collection |  |  |
| Days after symptoms offset | 21-28 | NA |
| Symptomatology during COVID-19 | Mild-Moderate | NA |
| Symptomatology at collection | Asymptomatic-Mild | Asymtomatic |
| PCR test | Positive (100%) | Negative (100%) |
| Rapid Immunochromatographic test | Positive (92%) | Positive 4% |
| Another ELISA test (IVD) | Positive (88%) | Positive 8% |
